# The Signature Microbiota Driving Rumen Function Shifts in Goat Kids Introduced Solid Diet Regimes

**DOI:** 10.1101/737775

**Authors:** Xiaokang Lv, Jianmin Chai, Qiyu Diao, Wenqin Huang, Yimin Zhuang, Naifeng Zhang

## Abstract

The feeding regime of early supplementary solid diet improved rumen development and ruminant production. However, the signature microbiota linking dietary regimes to rumen function shifts and hosts are still unclear. We analyzed the rumen microbiome and functions affected by supplementation of solid diet using a combination of machine learning algorithms. The volatile fatty acids (i.e., acetate, propionate and butyrate) fermented by microbes increased significantly in the supplementary solid diet groups. The predominant genera altered significantly from unclassified *Sphingobacteriaceae* (non-supplementary group) to *Prevotella* (supplementary solid diet groups) RandomForest classification model revealed signature microbiota for solid diet that positively correlated with macronutrient intake, and linearly increased with volatile fatty acids production. The nutrient specific bacteria for carbohydrate and protein were also identified. According to FishTaco analysis, a set of intersecting core species contributed with rumen function shifts by solid diet. The core community structures consisted of specific signature microbiota and their symbiotic partners are manipulated by extra nutrients from concentrate and/or forage, and then produce more volatile fatty acids to promote rumen development and functions eventually host development. Our study provides mechanism of microbiome governing by solid diet and highlights the signatures microbiota for animal health and production.

**Importance:** Small ruminants are essential protein sources for human, so keeping them health and increasing their production are important. The microbial communities resided in rumen play key roles to convert fiber resources to human food. Moreover, rumen physiology experience huge changes after birth, and understanding its microbiome roles could provide insights for other species. Recently, our studies and others have shown that diet changed rumen microbial composition and goat performance. In this study, we identified core community structures that were affected by diet and associated to the rumen development and goat production. This outcome could potentially allow us to select specific microbiome to improve rumen physiology and functions, maintain host health and benefit animal production. Therefore, it gives a significant clue that core microbiome manipulation by feeding strategies can increase animal products. To our knowledge, we firstly used FishTaco for determination of link between signatures abundances and rumen function shifts.

## Introduction

With the development of next generation sequencing, the roles of gut microbiome have been dramatically understood. The early life diet, especially introduction of solid diet, is an important driver in shaping long-term and adult gut microbiome profiles due to the novel alteration of diet components and macronutrient levels as well as gut anatomical development (1). Goat with rapid physiological changes (non-rumination, transition and rumination) could be proposed as an appropriate animal model for studying the gut microbial ecosystems development by early diet intervention and providing a means for prevention of metabolic diseases (2, 3). Young ruminants receiving only milk or fluid diet (milk replacer) have limited metabolic activity in the rumen epithelium and minimal absorption of volatile fatty acids (VFA) (4). Early supplementary feeding solid diet has been widely used in lamb production to improve rumen and body development since it can stimulate microbial proliferation and VFA production that initiates epithelial development (5). A solid concentrate diet (starter) containing high concentration of carbohydrate has been widely used to rear pre-weaned ruminants (4, 6, 7). Compared with breast milk-fed lambs, the community structure and composition of rumen microbiota of started-fed lambs tends to mature easily and quickly (8). Lin and their colleagues (3) analyzed rumen microbiota in lambs fed starter vs breast-milk. They found that acetate and butyrate increased in starter-feeding lambs, as well as increases of 5 genera including *Mitsuokella*, *Sharpea*, *Megasphaera*, *Dialiste*, and unclassified *Bifidobacteriaceae*. An extra alfalfa supplementation on the basis of concentrate diets improves rumen development to the next level. Previous studies reported that increases of growth performance and changes of ruminal microbiota during the pre- and post-weaning periods were found in lambs fed starter plus alfalfa compared with lambs fed fluid-diet and starter (9, 10). In addition, some studies have summarized the significant changes of microbiota in solid feeding regime and evenly calculated the correlation between macronutrient intake and rumen bacterial abundances. Wang et al. (11) found the correlation between bacterial genera in lambs rumen tissue and functional variables at d42. Yang et al. (10) sequenced rumen samples from *Hu* lambs fed milk replacer from d5 to d38 and supplied with solid diet (starter and alfalfa). They observed the effect of solid diet on microbial composition, and a set of taxon correlated with CP, NDF and body weight.

Until now, although these studies remarkably extend the effects of solid diet on the development of rumen functions and microbial communities in lambs, they mainly focus on the microbiota at weaning day (around d 40) or at genus level. Many key questions remain unclear. For example, does goat kids have similar pattern by solid diet since lambs and goats belong to different genus? What are the signature microbiota for supplementary regimes? How does the regime supplemented starter plus alfalfa affect rumen microbiota manipulation? How does the signature microbiota associated with other members in solid diet regime maintain equilibrium and improve function? To address these questions, a study that feeds goats with solid supplement to investigate microbiome and their association with experimental factor and rumen function using more machine leaning algorithms is urgent. RandomForest, an ensemble learning method for classification and regression, can be used to rank the importance of predictor variables in a regression or classification problem in a natural way (12). FishTaco, a computational framework for comprehensively computing taxon-level contribution to detected functional shifts and identifying key taxa, was introduced by (13). Network analysis that identify the microbial interaction allows us to characterize how the “core” microbiota impacts the overall composition and function (14).

Therefore, the objectives of this work was to assess the rumen fermentation, microbiome community and function shift influenced by supplementary solid diet fed until to d 60 (rumination phase). We addressed above questions by deeply analysis of microbial data with a combination of above three algorithms. Also, the correlation between phonotype, such as macronutrients intake and rumen fermentation parameters, and microbiome were identified. We observed that extra solid diet intake in early life could change the rumen microbial community structure towards maturated level by increasing signature microbiota qualitatively or quantitatively, and then fermentation environment and functions.

## Results

### Rumen fermentation parameters

The rumen fermentation parameters affected by the different dietary regimes were observed (Figure 1 & Table S3). The MRO group had greater concentration of NH_3_-N (*P*<0.05) compared with the MRC and MCA group, while the opposite pattern of ruminal microbial protein was found. The more concentration of total VFA, acetate, propionate, butyrate and valerate in supplementary solid diet regimes (MRC, MCA) as compared with MRO was observed (*P*<0.05), and except propionate and valerate, the acetate, butyrate and Total VFA were higher in MCA than MRC (*P*<0.05).

**Figure 1.**
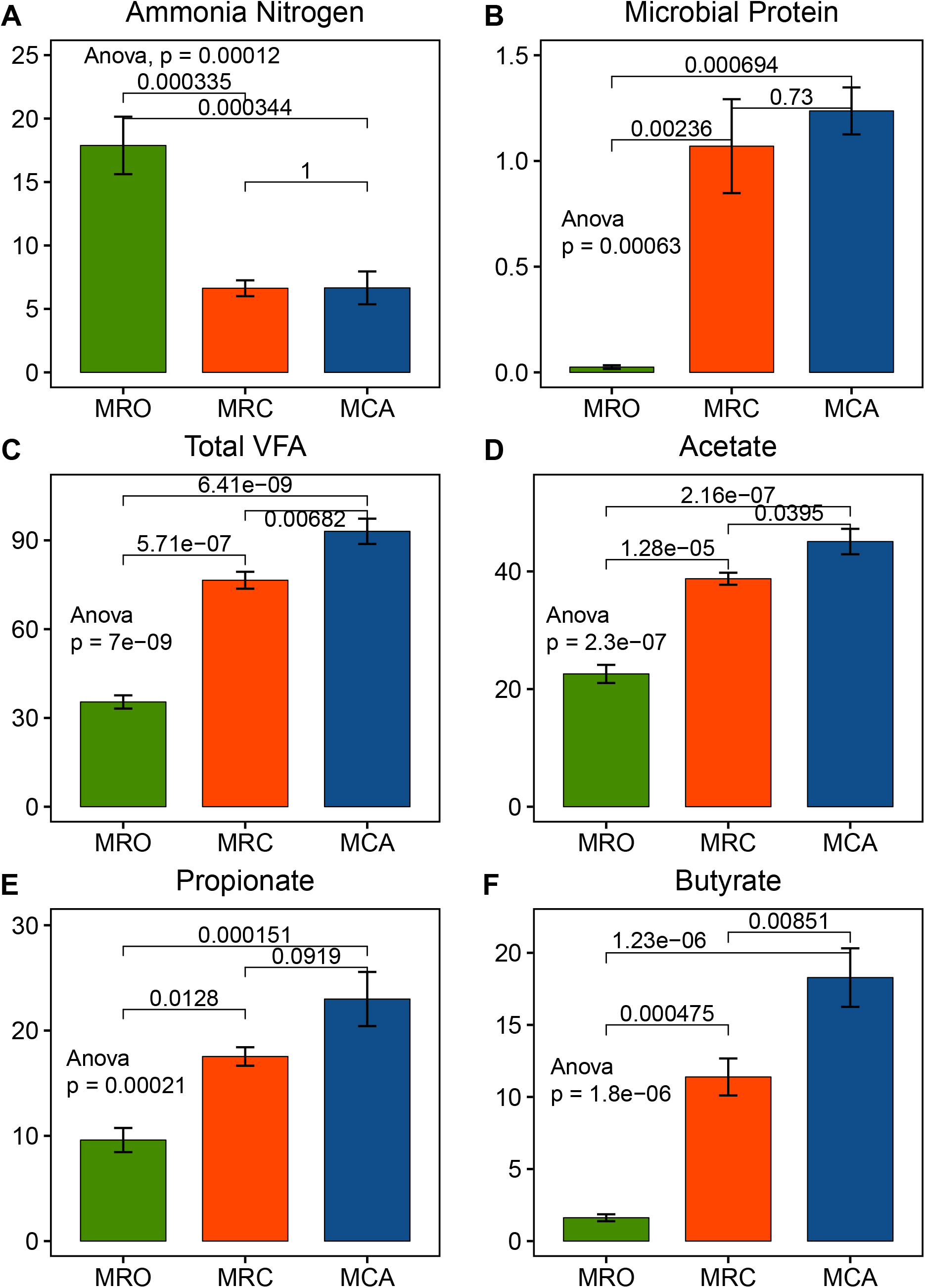
Effects of early supplementary solid diet on rumen fermentation parameters in goat kids The Anova test was used for significance calculation after detection of homogeneity of variance. After the global test was significant, a post-hoc analysis (Tukey’s HSD test) was performed to determine which group of the independent variable differ from each other group. High dietary nitrogen conversion ratio was found in MRC and MCO (*P*<0.05). The total VFA, acetate propionate and butyrate had the highest values in MCA (*P*<0.05), and they were significantly higher in MRC than in MRO (*P*<0.05). MRO=milk replacer only, MRC= milk replacer + concentrate, MCA= milk replacer + concentrate + alfalfa. VFA: Volatile fatty acids. Statistical significance was accepted at *P*<0.05.

Since the key factors of diets influenced goat rumen environment and development were intake of nutrient including CP, NFC and NDF, thus, correlation between nutrient intake and rumen fermentation parameters was performed (Table S4). Regression analysis confirmed that pH and NH_3_-N were negatively associated with average daily intake of CP, NFC and NDF, while rumen MCP and VFA (i.e., acetate, propionate, butyrate, and Total VFA) concentration had the strongly positive association with nutrient intake.

### The diversity and core bacteria in rumen microbiome

After quality control, filtering, and OTUs clustering steps, 64,1197 high quality sequencing reads across all samples and an average of 3,5622 sequence reads for each sample were generated. Firstly, we analyzed all the rumen content microbiome at community level. Although diversity (Shannon Index) was not different (p=0.372), significance of microbial richness was observed among MRO, MRC and MCA rumen samples (p=0.012) (Figure 2 A&B). The MRO rumen microbiota had significantly higher observed species than both MRC and MCA samples (p=0.045, p=0.005), and there was no difference between MRC and MCA (p=0.180). The observed species of rumen microbiome was negatively correlated with nutrient average daily intake including CP (r=-0.65, p=0.003), NFC(r=-0.73, p=0.001) and NDF (r=-0.74, p=0.0003) (Table S5). Negative association between microbial richness and MCP and VFA including acetate, propionate, butyrate, valerate and Total VFA was also observed. Regarding the beta diversity measurements, significant cluster in community structure among 3 regimes were detected (Weighted Unifrac ANOSIM, R=0.68, P<0.05; UnWeighted Unifrac ANOSIM, R=0.69, P=0.001). The MRO formed a distinct cluster (green dots) on the left side, while the MRC and MCA were closely clustered (red and blue dots) on the right side of PCoA plot (Figure 2 C&D).

**Figure 2.**
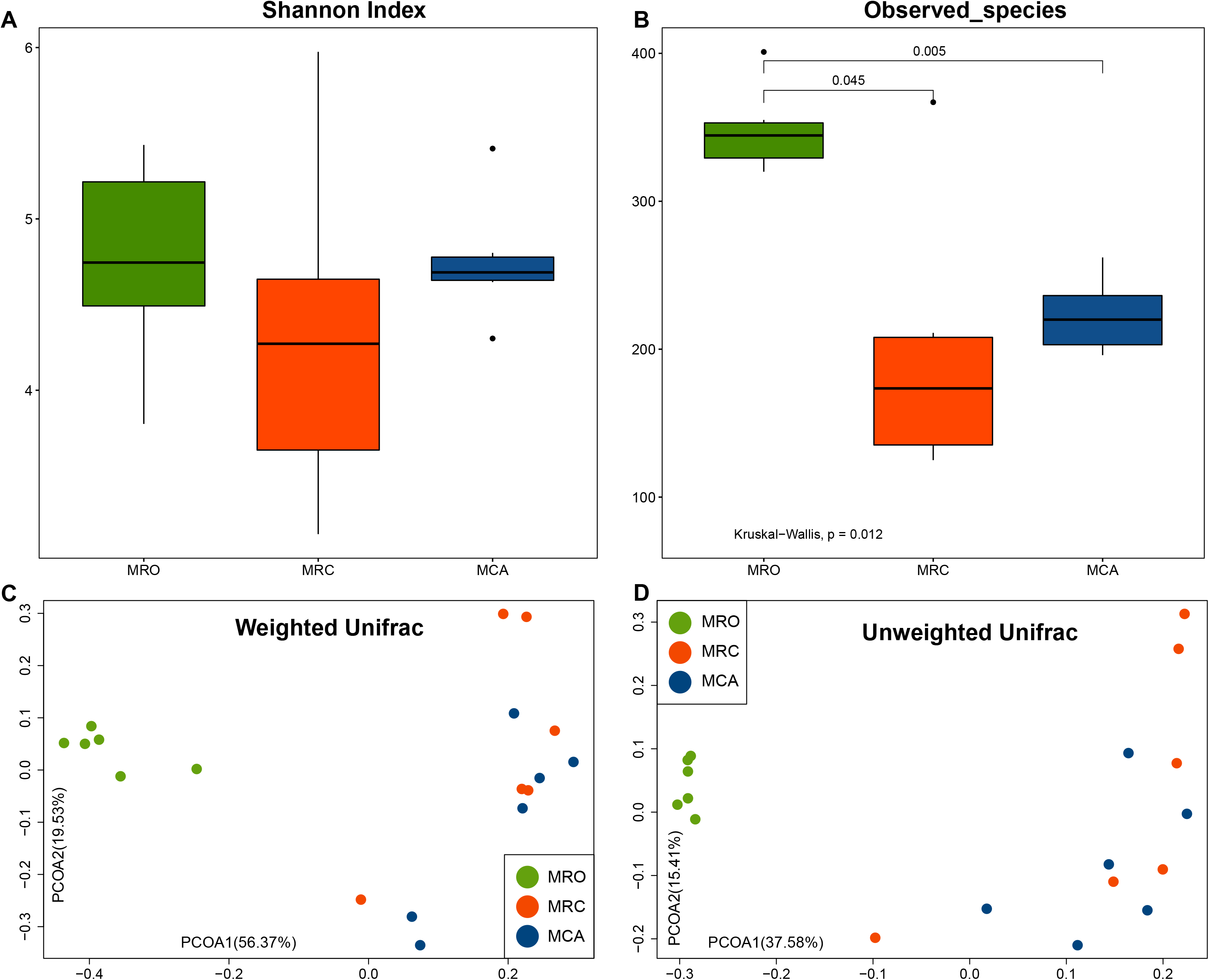
The early supplementary solid diet impacted on Alpha and Beta diversity of rumen microbiome in goat kids. (A-B) The Shannon Index and Observed species. Alpha diversity of the rumen microbial data was tested using Kruskal-Wallis test and a post-hoc Dunn Kruskal-Wallis multiple comparison with Bonferroni method for p value correction. Principal coordinate analysis (PCoA) of the community membership based on the weighted (C) and unweighted (D) UniFrac distance, with the green cycles as MRO, red cycles as MRC and blue cycles as MCA. Although diversity (Shannon index) was not different (p=0.372), significance of microbial richness was observed among MRO, MRC and MCA rumen samples (p=0.012). Significances in community structure among 3 group were detected (Weighted Unifrac ANOSIM, R=0.68, *P*<0.05; UnWeighted Unifrac ANOSIM, R=0.69, *P*=0.001). The MRO formed a distinct cluster in on the left side, while the MRC and MCA were closely clustered on the right side of PCoA plot. MRO=milk replacer; MRC= milk replacer + concentrate; MCA= milk replacer + concentrate + alfalfa; ANOSIM: Analysis of similarity

We next examined the rumen core microbiome among three treatments. At genus level, a total of 152 genera were observed, and *Prevotella* followed by unclassified *Prevotellaceae*, unclassified *Sphingobacteriaceae* and unclassified *Bacteroidetes* accounted for 63.29% of the total sequences were the predominant genera with abundance over 5% across all samples(Figure S1). The top genera in MRO were unclassified *Sphingobacteriaceae* (30.32%), unclassified *Prevotellaceae* (16.92%), unclassified *Bacteroidetes* (11.77%) and *Prevotella* (8.91%). In MRC, *Prevotella* (56.02%) were the predominant bacteria, followed by *Roseburia* (4.49%), unclassified *Prevotellaceae* (4.29%), *Selenomonas* (3.82%) and unclassified *Lachnospiraceae* (3.73%). However, in MCA, the abundance of the predominant genus *Prevotella* (44.02%) decreased compared with MRC, and other dominant genera were unclassified *Prevotellaceae* (11.63%), *Fibrobacter* (7.01%), *Treponema* (5.35%), *Succinivibrio* (4.74%) and unclassified *Lachnospiraceae* (4.50%)

At OTU level, there were 281 OTUs that were significantly different between 3 groups (Table S6), and 16 taxa in top 30 were significant. The top 30 most abundant bacterial taxa accounting 57.77% of all reads are displayed on stacked bar charts (Figure 3). Among top 30 OTUs, 14 belong to genus *Prevotella*, and 4 was owned by genus unclassified *Prevotellaceae*. The OTUs belong to unclassified *Sphingobacteriaceae* (OTU1 and OTU5), unclassified *Prevotellaceae* (OTU4 and OTU30) and OTU24 *Cloacibacillus* were greater in MRO. The OTUs affiliated with *Prevotella* (OTU2, OTU6, OTU13,) in top 30 had higher abundance in MRC and MCA. The MRC were abundant with OTU10-*Roseburia* OTU20-*Olsenella* and OTU21-*Prevotella*. The bacteria belong to genera of *Prevotella* (OTU6 and OTU13) *Succinivibrio* (OTU9), unclassified *Prevotellaceae* (OTU15), *Succiniclasticum* (OTU22) had the highest relative abundance in MCA.

**Figure 3.**
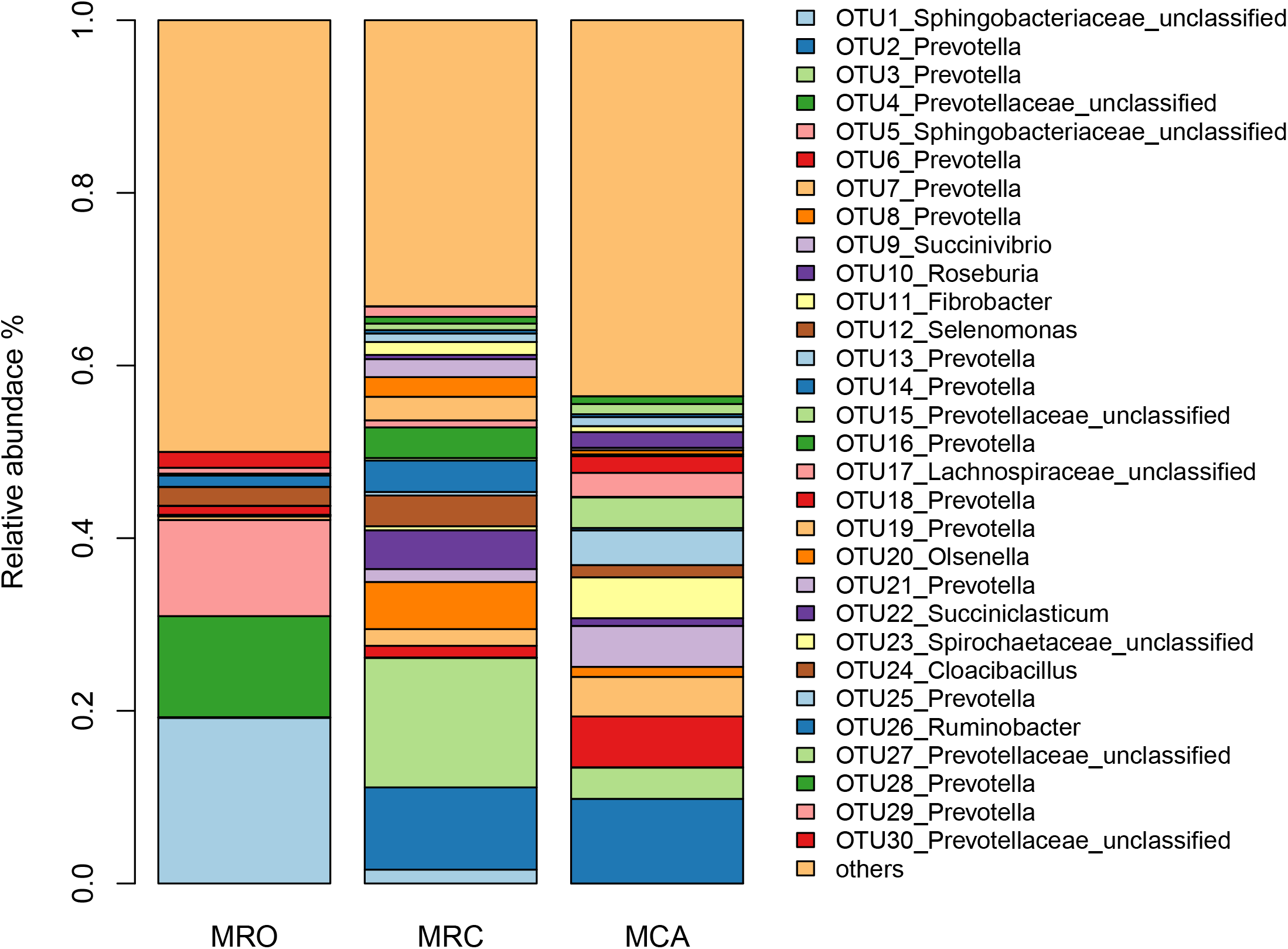
The top 30 OTUs in 3 supplementary regimes. Each bar shows the average relative abundance of MRO, MRC and MCA. Each color represents the relative abundance of a bacterial taxon on the stacked bar chart. MRO=milk replacer, MRC= milk replacer + concentrate, MCA= milk replacer + concentrate + alfalfa.

### The signature microbiota differentiating MRO, MRC and MCA supplementary regimes

To identify the rumen important microbiome that differentiate MRO, MRC and MCA, we performed an updated RandomForest classification model to differentiate these 3 supplementary regimes. The regimes-associated bacterial features were listed based on their MDA and the representatively selected microbiota were presented in Figure 4. All 3 groups were analyzed together, and optimal features with an AUC (area under the curve) of 1.00 (specificity 1.00, sensitivity 1.00) were selected from AUCRF model (Table S7; Figure S2). High AUC (0.931) was still observed at 50^th^ feature suggesting those signatures being able to accurately predict whether goats was fed concentrate plus alfalfa. Among top 50 features, only 3 core OTUs such as OTU5 (unclassified *Sphingobacteriaceae*), OTU24 (*Cloacibacillus*) and OTU6 (*Prevotella*) were identified as regime-associated bacteria (Figure 4). Forty of top 50 bacteria were more abundant in MRO. OTU5 associated with MRO predominant genus had more relative abundance and prevalence (11.13%; 6/6) as compared with MRC (0.03%, 2/6) and MCA (0.04%, 2/6). OTU24 as qualitative signatures had more abundance 2.16% in MRO. Other species associated with *Prevotella* that was enriched genus in solid diet groups were also found more abundant in MRO, including OTU119, OTU42 and OTU60. For MCA microbiome, OTU6 and OTU104 affiliated with predominant *Prevotella* increased. We observed the relative abundances of OTU6 was 0.01% 1.35% and 5.89% in MRO, MRC and MCA (prevalence 2/6, 6/6 and 6/6). OTU87 (*Butyrivibrio*) and OTU83 (unclassified *Bacteroidales*) were significantly enriched in MCA and extremely low abundance in MRO and/or MRC. Similar patterns could be found in other MCA predictors such as OTU93 and OTU74 (*Treponema*), OTU539 (unclassified *Clostridiales*), OTU396 (unclassified *Proteobacteria*), OTU221 (*Pseudobutyrivibrio*) and OTU110 (unclassified *Prevotellaceae*) (Figure S3-1 & S3-2).

**Figure 4.**
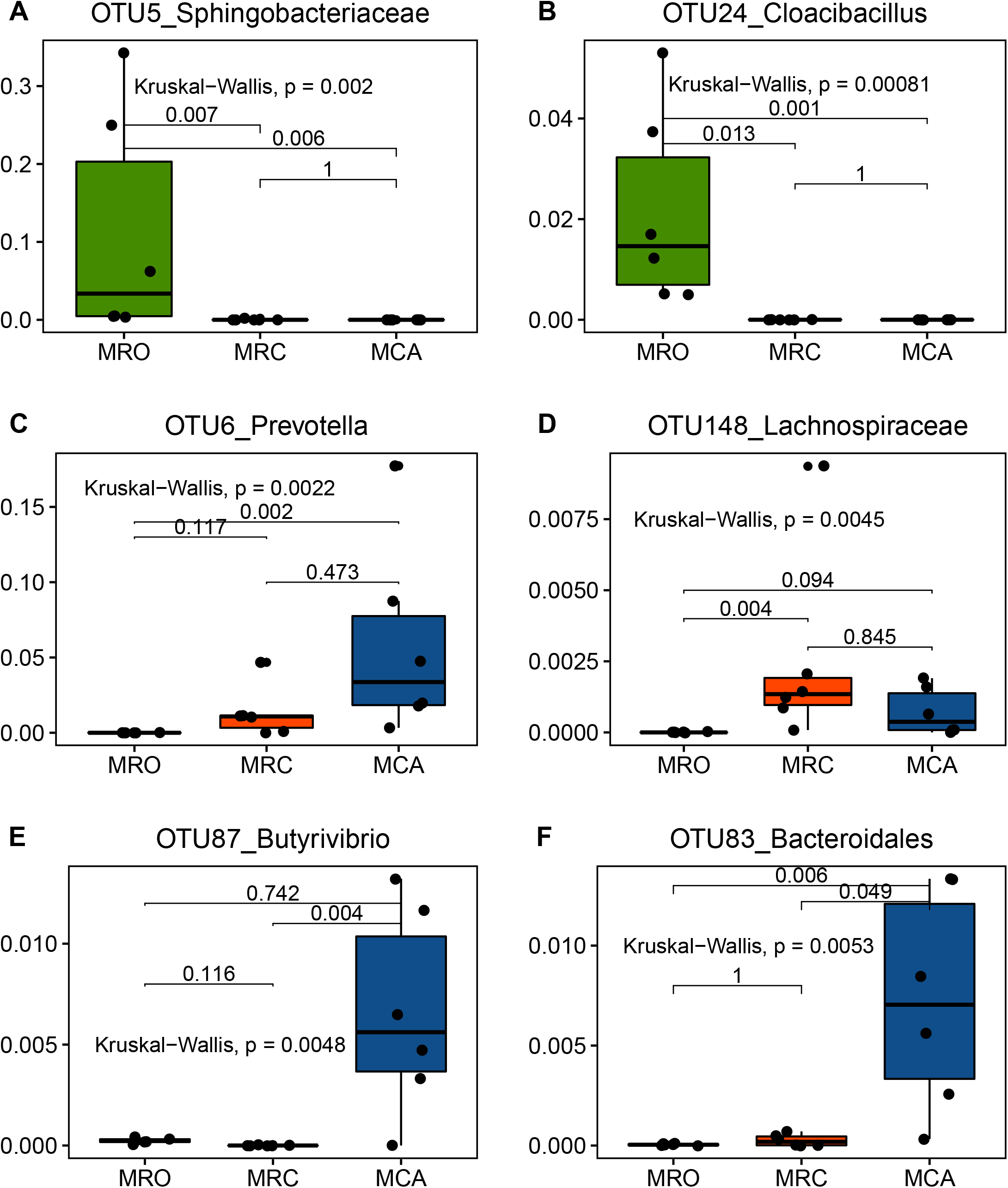
The highlight signature microbiota identified by AUCRF for differentiating MRO, MRC and MCA All the OTUs abundances were tested using Kruskal–Wallis test and a post-hoc Dunn Kruskal-Wallis multiple comparison with Bonferroni method for p value correction. The black dots within each bar were values from individual animal, and the black lines within each bar represented the medians. MRO=milk replacer; MRC= milk replacer + concentrate; MCA= milk replacer + concentrate + alfalfa; AUCRF: RandomForest based on optimizing the area-under-the receiver operator characteristic curve (AUC);

Then, we performed pair wise AUCRF comparisons to validate these predictors. The results confirmed that most of the classified biomarkers could also be listed (Figure S4-S6). Moreover, MRC were enriched with OTU148 (unclassified *Lachnospiraceae*) and OTU114 (*Megasphaera*) compared with MRO, whereas it had more abundance of OTU643 (*Neisseria*), OTU177 (*Campylobacter*) and OTU314 (*Blautia*) as compared with MCA.

### Phenotypes and rumen microbiota

To find the relationship between rumen microbiota with major nutrients of diet for better understanding how supplementary feeding regimes influenced microbial communities. Firstly, we performed RandomForest regression model by using CP, NFC and NDF intake as outcomes and all taxa as independent variables. Then, the Pearson correlations were calculated between selected top 50 bacterial abundances and dietary CP, NFC and NDF intake respectively (Figure 5). On other hand, the impacts of alteration of rumen microbiota on rumen VFA were also estimated using same approaches.

**Figure 5.**
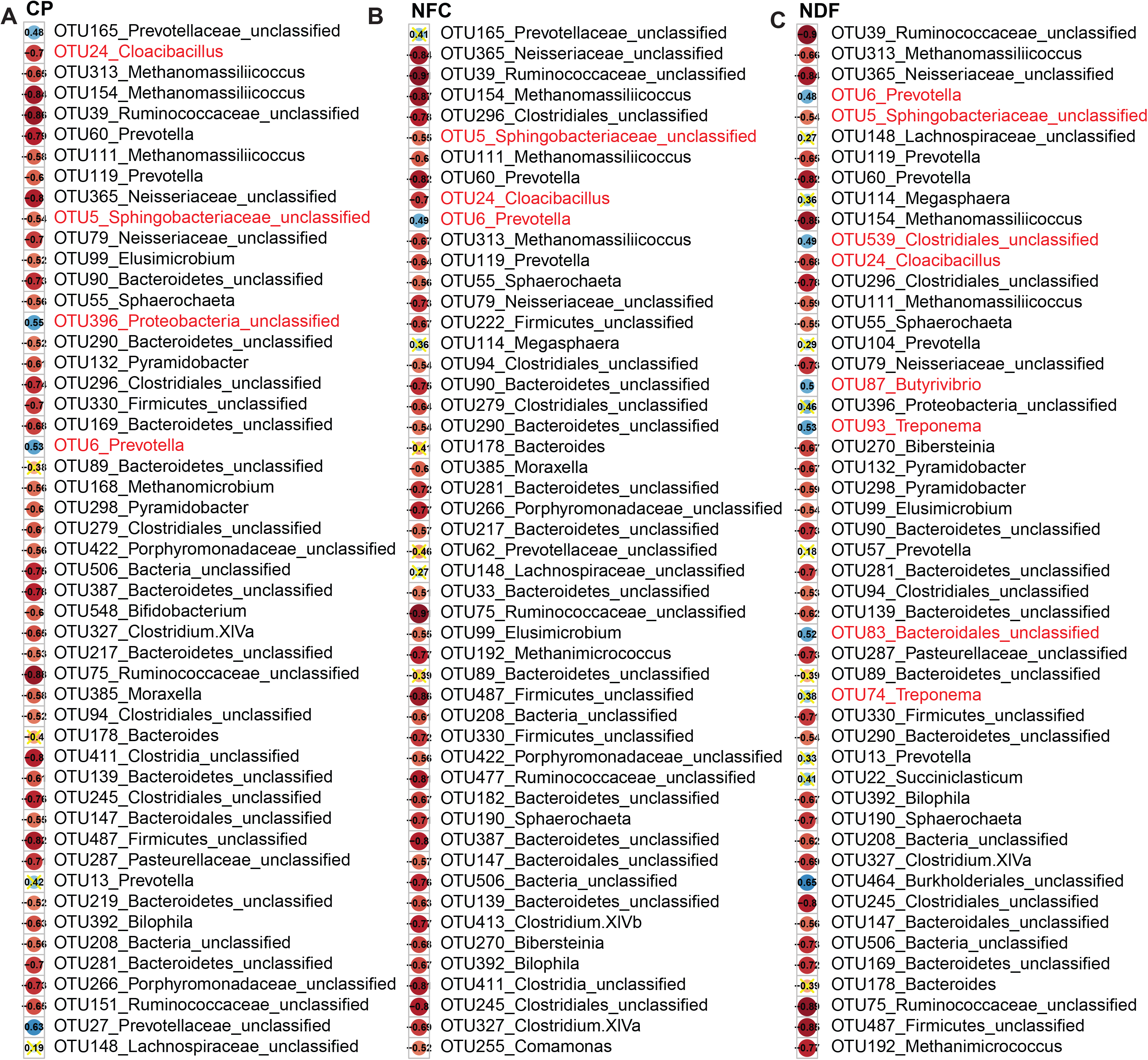
Correlation analysis between nutrient (CP, NFC and NDF) intake and rumen microbes in goat kids. We performed the RandomForest regression model across all samples between dietary average daily CP, NFC and NDF intake and all the genera with high prediction accuracy (Table S6). Then, using the abundances of top 50 features to calculate the pearson correlation with intake of CP, NFC and NDF was carried out. We consider p<0.05 as a significant correlation and yellow crosses indicated non-significant. The Pearson coefficients were labeled black values inside cycles, red dots represented negative correlation, and blue dots indicated a positive correlation. The bacteria from up to bottom followed the descending order of mean square error. CP: Crude protein average daily intake; NDF: Neutral detergent fibers average daily intake; NFC: Non-fibrous carbohydrates average daily intake.

The rumen microbiota had high prediction accuracy (>73%) to explain nutrients intake (Table S8). Among CP, NFC and NDF, 31 shared bacteria were observed, and 30 of 31 were the predictors identified by RandomForest classification model. In these shared bacteria, 27 as MRO-associated predictors had negative correlation with intake of CP, NFC and NDF, such as OTU5, OTU24. For other 3 shared features, OTU327 (*Clostridium XlVa*) negatively correlated with intake of CP, NFC and NDF, OTU148 (unclassified *Lachnospiraceae*) had no correlation (p>0.05), and OTU6 (*Prevotella*) was positively and moderately correlated with them (r=0.53, 0.48, 0.49, p=0.023, 0.043, 0.041). Regarding to CP and NFC intake, OTU165 (unclassified *Prevotellaceae*) was the shared OTUs increasing abundances. The OTU396 (unclassified *Proteobacteria*) and OTU27 (unclassified *Prevotellaceae*) were specifically and positively correlated with CP intake (r=0.55, 0.63, p=0.019, 0.005). When NDF intake was observed, the abundances of its associated microbiota went up. For example, OTU464 (unclassified *Burkholderiales*) increased with more NDF intake (r=0.65, p=0.003), while others identified as predictors for MCA (i.e., OTU87, OTU83, OTU93, and OTU539) also linearly increased abundance with increase of NDF intake (r=0.53, 0.52, 0.49, 0.50 and 0.48). Interestingly, OTU74 (*Treponema*) identified as MCA signature had no significant association with NDF intake (r=0.38, p=0.124). Moreover, the OTU396 and core significant *Succiniclasticum* (OTU22) tended to moderately correlate with NDF intake (r=0.46, p=0.051; r=0.41, p=0.09).

Although the regression prediction of MCP and NH_3_-N was not high (50.97% and 44.94%), OTU6 and OTU27 correlated with CP were associated with NH_3_-N, and other regime-associated signature (OTU148) significantly correlated with rumen nitrogen indexes (File S1). Moreover, OTU152, OTU268 and OTU322 had significant correlation with MCP. The rumen microbiota also had accurate prediction for VFA concentration (Table S8). Shared OTUs were also found in the list of between RandomForest classification and VFA regression (i.e., 39 acetate, 17 propionate, 24 butyrate, 25 valerate, 36 Total VFA). Those shared OTUs were most of MRO-associated signatures and negatively correlated with VFA. For the bacteria positively correlated with Total VFA, they were also observed within 1 or 2 of acetate, propionate or butyrate regression models, for example, OTU6 within Total VFA and butyrate; OTU396 within Total VFA and propionate and butyrate. Considered the major VFA (acetate, propionate and butyrate), OTU83 was the only common microbes were correlated positively with all of them (r=0.63, 0.54, 0.56; p=0.005, 0.020, 0.017). When increasing acetate was observed, the abundances of OTU122 (*Ruminobacter*), OTU143 (*Fibrobacter*) and OTU204 (unclassified *Bacteroidetes*) tends to increase. Regarding to propionate, positive correlation were found in the bacteria, such as OTU13 (*Prevotella*), OTU93 (*Treponema*), OTU165 (unclassified *Prevotellaceae*), OTU258 (*Olsenella*), OTU120 (Megasphaera), OTU532 (unclassified *Bacteroidetes*), OTU322 (*Allisonella*), OTU604 (*Eubacterium*) and OTU530 (*Mitsuokella*). The butyrate-associated bacteria were OTU6, OTU13, OTU539, OTU15 (unclassified *Prevotellaceae*), OTU17 (unclassified *Lachnospiraceae*), OTU114 (*Megasphaera*) and OTU205 (unclassified *Firmicutes*). Notably, higher ensemble prediction score (70%) in valerate regression indicated that rumen microbiota explained it better as well. When valerate increased, unclassified *Lachnospiraceae* (OTU148 and OTU391), *Olsenella* (OTU20, OTU258), *Megasphaera* (OTU114, OTU120 and OTU173), unclassified *Clostridiales* (OTU311), *Mitsuokella* (OTU152), unclassified *Bacteria* (OTU52), *Prevotella* (OTU186) and unclassified *Porphyromonadaceae* (OTU47) linearly increased.

### Rumen microbiota driving function shifts

To predict how rumen microbiota associate with solid diet supplementary regimes, PICRUSt based on OTUs’ level was used to predict the abundances of functional categories the KEGG. In the level 3, nutrient pathways were the most popular in this study (Figure S7). Many bacterial genes in all 3 groups could potentially trigger pathway function of same nutrient metabolism, but different treatments participated different reaction modules. For example, carbohydrate metabolism found in all groups had the specific reaction of pyruvate metabolism and citrate cycle in MRO; Fructose, mannose Starch and sucrose metabolism in MRC; and glyoxylate and dicarboxylate metabolism in MCA. Moreover, some cellular process pathways were found in goat supplied with solid diet. MRC was enriched membrane transport (ABC transporters) and Insulin signaling pathway. The pathways of transcription factors and machinery were found in MRC and MCA.

The FishTaco was performed to identify the corresponding microbiota driving the functional shifts between supplementary regimes. There were no differences of normalized abundance of functions between MRC as control and MCA as case based on Wilcoxon rank-sum test. When MRO as control and case defined as MRC and MCA separately, 31 and 37 significant pathways were found (File S2-S3). Notably, 21 shared functions were observed between 2 comparisons, including metabolism of nutrient (lipid, amino acid, carbohydrate, vitamin, peptidoglycan, terpenoids and polyketides), and the pathway of endocrine system and cellular processes. (Figure 6 & S8-S9). To better understand the driver OTUs function, we get all sequences identifier with the highest scores on NCBI BALSTN database (File S4). Across all significant functions enriched in MRC, a set of *Prevotella* bacteria including OTU3 (*Prevotella copri DSM*), OTU2 (*Prevotella brevis strain GA33*), OTU14 (*Prevotella histicola*) and OTU16 (*Prevotella ruminicola*) (occurrence 100%, 83.3%, 60% and 13.3%) were the main drivers (Figure S8-S9). While in MCA, the function shifts were driven by a convoluted outcome of *Fibrobacter* and *Prevotella* including OTU11 (*Fibrobacter succinogenes*), OTU2, OTU3, OTU7 (*Prevotella ruminicola*) and OTU13 (*Prevotella brevis strain GA33*) (their occurrence 100%, 100%, 100%, 30.3% and 15.2%). Although selenocompound metabolism pathway was enriched in MRC and MCA compared with MRO, the set of OTUs drove this enrichment of 2 regimes and the level of contribution of each specie differed, with OTU2, OTU3 and OTU14 driving the shift in MRC and a set of OTU2, OTU3, OTU11, and OTU13 in MCA (Figure 6). These enrichments were attenuated by greatly different bacteria in MRC (OTU20, OTU16, OTU8, OTU10, OTU12 and OTU14) and MCA (OTU15, OTU6, OTU7, OTU17, OTU22 and OTU9). Other highlight pathways such as lipid and carbohydrate metabolism, Biosynthesis of unsaturated fatty acids and transporters had similar pattern. In addition, the MRO enriched microbiota including unclassified *Sphingobacteriaceae* (OTU1 and OTU5 *Olivibacter sitiensis*) and *Cloacibacillus* (OTU24 *Cloacibacillus porcorum*) were strongly depleted by solid diets.

**Figure 6.**
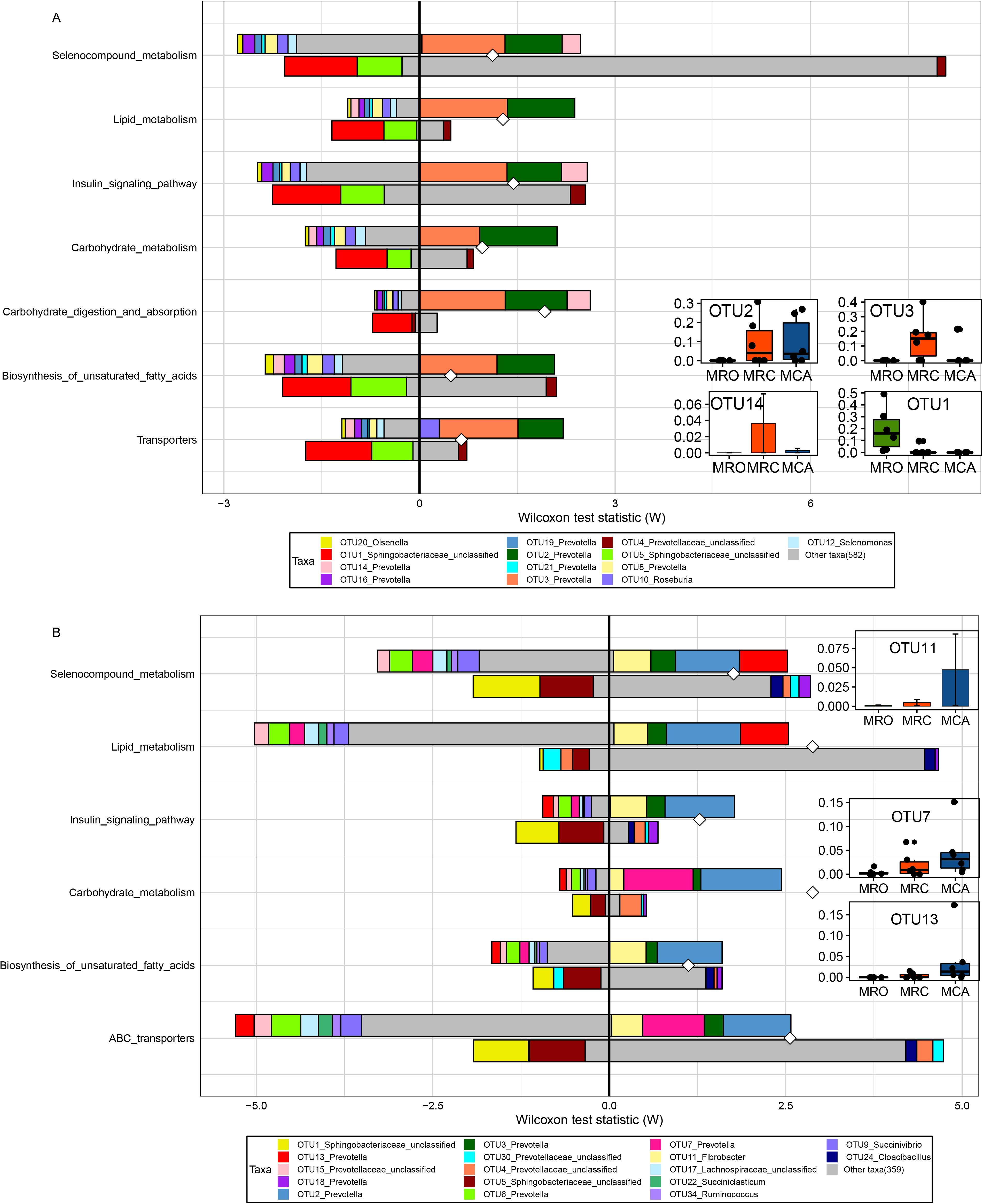
Comparing taxon-level contribution profiles of functional shifts: A: MRO control and MRC case, B: MRO control and MCA case. Taxon-level shift contribution profiles for case-associated functional modules by FishTaco. The horizontal axis represents rank and statistic scores, and the vertical axis represents related pathways. For each functional pathway, the bar on the top-right of Y axis represents case-associated bacteria driving the enrichment in the functional module; the bar on the top-left of Y axis indicates case-associated bacteria attenuating functional shift; the bar on the bottom-right of Y axis represents bacteria depleted in control driving functional shift; the bar on the bottom-left of Y axis shows bacteria depleted in control attenuating functional shift. White diamonds represent bacterial-based functional shift scores. The abundances of main drivers were displayed on the right side. OTU2 and OTU3, the shared drivers of enrichments of MRC and MCA were abundant in solid diet regimes. OTU1 enriched in MRO was strongly depleted by solid diets. OTU11, OTU7 and OTU13, driving mainly MCA function shifts, increased abundance with supplementation of solid diet. FishTaco:Functional Shifts’ Taxonomic Contributors; MRO=milk replacer; MRC= milk replacer + concentrate, MCA= milk replacer + concentrate + alfalfa;

### Network analysis of regime associated microbiota

Network analysis revealed core sun-community structure within communities that consisted of a set of bacteria associated with the phenotypes and rumen functions in the supplementary regimes. We detected respectively 4, 7 and 8 main subnetworks in MRO, MRC and MCA (Figure 7). The species that were observed as regime-associated features and identified as functions drivers formed the main subnetwork. In MRO, the predictors, OTU60 (violet cluster), OTU42 and OTU111 (green cluster), OTU99, OTU79, OTU55 (yellow cluster) and OTU33, OTU94, OTU89 (pink cluster) formed the main subnetwork, showing significant correlations with a large number of other members of MRO community. For MRC rumen microbiota, OTU2, OTU6 OTU16 within palegreen cluster, OTU3 within pink cluster and OTU14 within yellow cluster, as dominate species associated with other members, consisted of the main subnetworks. Within MCA, OTU104, OTU11, OTU2, OTU6, OTU13, OTU87, OTU74, OTU83 and OTU3 recognized as main drivers or signatures were the main members of subnetworks. Their partners interacted with these microbiota may associate with fermentation, such as OTU7 and OTU27 in MCA. Moreover, OTU79 (*Snodgrassella alvi*) and OTU99 (*Elusimicrobium minutum*) as hub nodes in MRO connected yellow and blue cluster, whereas OTU87 (*Butyrivibrio hungatei*), OTU15 (*Metaprevotella massiliensis*) and OTU31 (*Fibrobacter succinogenes subsp. Elongates*) served as a bridge to link three clusters.

**Figure 7.**
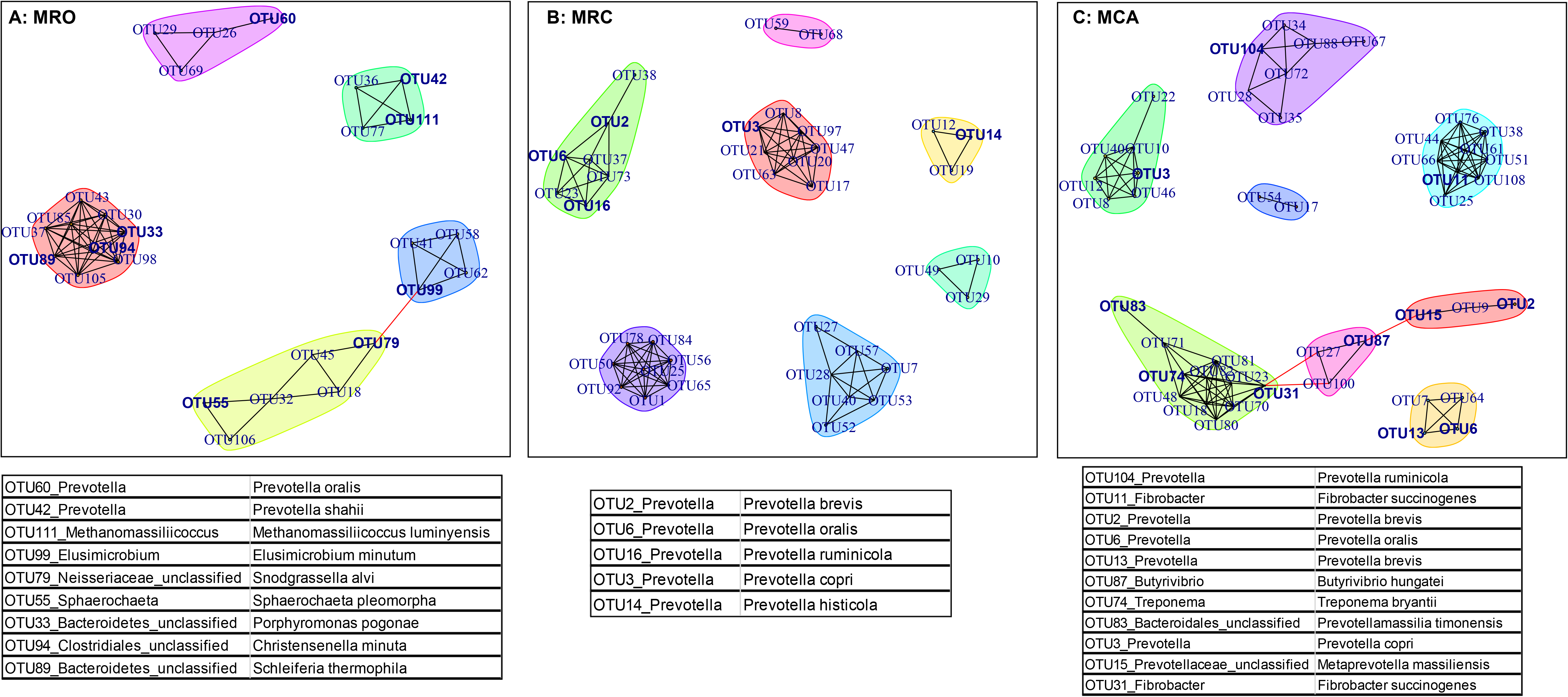
Network analysis of the interactions between bacterial taxa at MRO (A), MRC (B) and MCA (C). The OTUs accounting for >0.5% of the total sequences were selected to network analysis. Each node denotes a particular OTU within the network and each line (edge) a significant co-efficiency relationship (Pearson rank correlation coefficient >0.6 or <−0.6. The table under corresponding figures contained the highlight OTUs and their sequence identifiers with the highest scores from NCBI database (other OTUs identifiers were shown in File S4). MRO=milk replacer; MRC= milk replacer + concentrate; MCA= milk replacer + concentrate + alfalfa; legend for supplemental materials

## Discussion

Early supplementary feeding with solid diet has a positive impact on rumen development by influencing rumen microbial population and composition, environment alteration and functional achievement. However, lack of information of microbial predictors for supplementary regimes leads to unclear mechanism of manipulation of rumen microbiota and function shifts. This study confirmed that rumen VFA especially acetate, propionate and butyrate increased significantly with the supplementation of solid diet, and promoted rumen weight and functions. The predominate genera changed from unclassified *Sphingobacteriaceae* to *Prevotella* when goat kids were supplied solid diet. The signature microbiota in corresponding feeding regimes significantly correlated with phonotypes such as major nutrients intake and VFA concentration. For example, the biomarkers for MCA (OTU6, OTU87, OTU83, OTU93 and OTU539) were positively correlated with NDF intake and VFA production. The improved rumen function in goats supplied solid diet were caused by the core bacteria, such as OTU3 (*Prevotella copri DSM*), OTU2 (*Prevotella brevis strain GA33*), OTU14 (*Prevotella histicola*) and OTU11 (*Fibrobacter succinogenes*). All these signatures and/or core microbiome formed main sub-communities to response solid diet feeding and drive function shifts.

The VFAs that are products of the fermentation of diets are essential to the rumen papillae development and nutrient source for host requirements (15). In ruminants, VFA produced in the rumen meets 70–80% of the energy requirement for the rumen epithelia, and 50–70% of the energy requirement for the body (16). In this study, rumen microbial proteins and VFA concentration increased in supplementation of solid diet. Other studies also revealed that early starter and alfalfa consumption facilitated rumen development and changed the pattern of ruminal fermentation (9, 11, 17). Moreover, we found the rumen microbial proteins and VFAs were positively correlated with intake of CP, NDF and NFC. Previous study reported that ruminal NH_3_-N increased linearly in response to increasing dietary CP (18). This study confirmed that microbiota in goats fed solid diet had more strong ability for biosynthesis of microbial proteins and VFA. Moreover, except the physical stimulation from solid diet, the chemical effects of nutrient intake may be another reason leading to increase of VFA. Therefore, early supplementation of solid diet leading to high nutrient intake increases rumen VFA production and nitrogen utilization efficiency, which reflects that microbiome experienced solid diets had a strong ability to utilize nutrient.

In pace with the change of rumen environment, this study also observed that the membership and structure of microbiota also altered by supplied with concentrate or forage compared with only fluid diet groups. Significant lower alpha diversity in starter feed-lambs and distinct beta diversity between starter feed- and breast milk-fed lambs were also reported (11). High bacterial richness in fluid diet groups might be temporary phenomenon at d60. Others confirmed that rumen microbiota at d70 had a lower richness compared with it at d42 (8). As we know, rumen is in rumination phase after 8 weeks and in transition phase in 3-8 weeks. In this study, compared with MRO, rumen microbiota in goats supplied with solid diet at d60 may have a more mature rumen function and stable microbiome structure at the same age. Another reason for reduction of richness in solid feeding regimes might be due to high concentration of VFA and low pH (19). In addition, the rumen microbiota in solid supplementary regimes had similar alpha and beta diversity. The similar pattern also could be observed in rumen fermentation parameters. This might be due to less feed intake of alfalfa and similar concentrate intake. In animal trial, the MCA goats had *ad libitum* access to concentrate and alfalfa pellets in separately troughs. Based on feed intake results, the goats preferred concentrate. Thus, future studies have to increase roughage intake for its effects on rumen microbiota or detect the microbiome after weaned milk replacer.

RandomForest is a powerful tool capable of delivering performance that is among the most accurate methods (20). It has been widely used in human microbiome studies to find the signatures for disease or healthy (21, 22). This study identified important signatures from 838 OTUs using RandomForest, which could provide more effective and accuracy information how diet supplementary regimes affected microbial composition. A higher AUC value (AUC=1.00) indicates the features are more efficiently classified. RandomForest not only gave an important score to the significant species but also find the accurate bacteria for experimental factors. For example, the low abundances of OTU87, OTU83, OTU93 and OTU539 were identified as the important predictor for regimes, which indicated that low abundance bacteria may also paly critical roles in function drifts. Therefore, previous literature (3, 10) only focused on the genera with significantly different abundances may not provide the best conclusion. RandomForest regression is useful and robust method for correlation applications because of its ability of automatically producing accuracy estimation and measuring the variable importance. Using it to select microbiota with high important scores would be a better and corrected method for finding precise signatures. The percent explained variance is a measure of how well out-of-bag predictions explain the target variance of the training set. High % explained variance (over 70%) in this study were found in CP, NFC and NDF model, which indicated that those top microbiota were more important to the responders. It helps to figure out the relationship between specific nutrient and microbiota. In addition, using FishTaco to link the microbiota abundances and rumen function shift caused by supplement of solid diet is a creative attempt (13). We used it to integrate comprehensively the significant species and function shifts. Compared with the original PICRUSt result, there was an improved result of significant pathways. Its process rely on a permutation-based approach, carefully designed normalization and scaling schemes to preserve overall community taxonomic characteristics and to account not only for variation induced by each bacteria but also for the way this variation correlated with community-wide context. Our results identified that a set of core bacteria were the main taxon drivers since low taxonomic abundance profiles were filtered and normalized. Finally, we observed subnetworks formed by these signature microbiota and their partners. In a word, more algorithms gave insights how microbiota had impacts on function shifts.

Rumen microbiota degrade fibers polysaccharides and proteins in diet, and yield VFAs and microbial proteins, which offer nutrients to meet the host’s requirement for maintenance and growth (23, 24). Based on these machine leaning algorithms, we analyzed microbiome to link supplementary regime to alteration of rumen environment and function. Supplementary solid diet altered the core microbiota from unclassified *Sphingobacteriaceae* to *Prevotella*. The representative OTU5 associated with unclassified *Sphingobacteriaceae* as MRO predictors were negatively with macronutrients intake and VFA production. OTU5 classified as *Olivibacter sitiensis* function is not clear, but it decreased with pH reduction when intake high concentrate diet (25). The species affiliated with *Prevotella* (OTU2, OTU3, OTU6 and OTU13) in top 30 increased in solid supplementary regimes. Other studies also reported that the abundances of the genus *Prevotella* that was predominent in starter-fed lambs positively correlated with acetate, propionate and urea nitrogen concentration (3, 8, 11). This genus is good at utilization of proteins and carbohydrates (either fiber- or non-fiber-carbohydrate) (26). Importantly, OTU6 (*Prevotella oralis*) was identified as signature species for solid diets, correlated with macronutrient intake and butyrate concentration. Another microbiota, OTU13 (*Prevotella brevis strain GA33*), was not classified as predictors for MCA, but we observed it had high abundance in MCA, positive association with propionate and butyrate, driving function shifts and interaction with other core microbiome. Therefore, increased abundance of these 2 species represented as *Prevotella* in rumen accessed solid diets promoted the improvement of rumen digestibility and function by yielding more VFA products. OTU2 (*Prevotella brevis*) and OTU3 (*Prevotella copri*) were the main drivers for functions shifts by solid diet. De Filippis et al. detected distinct strains of *Prevotella copri* by metagenome studies and showed that fiber-rich diets were linked to these strains with improved potential for carbohydrate catabolism (27). Broadly, introduction of solid fiber-rich diet to goat before weaning, *Prevotella* proliferated, which was also observed in other large domestic animals (28). They could be used as potential microbiota to utilize solid diet, maintain rumen community balance and prevent metabolic disease caused by dysbiosis. Another core genus increased in both solid diet regimes was unclassified *Lachnospiraceae*. The family *Lachnospiraceae* contains many known plant degrading species and most of the butyrate-producers (29). In our results, OTU148-*Lachnospiraceae* (*Kineothrix alysoides*) enriched in MRC was significant associated with NH_3_-N and valerate, although it was identified by regression model for nutrients intake and fermentation parameters. Regarding to others dominant bacteria, whereas *Roseburia* and *Selenomonas* specifically increased in concentrate diet regime. The abundances of *Fibrobacter*, *Treponema* and *Succinivibrio* arose became greater in extra supplementation of alfalfa. *Succinivibrio*, a saccharolytic bacteria, can yield acetate and lactate (30). The OTUs associated with these genera were not observed well in our study. For example, OTU10-*Roseburia* and OTU9-*Succinivibrio* formed main structure with other members; OTU74 affiliated with *Treponema* predicted MCA well but no relation with phonotypes; and OTU11-*Fibrobacter* drove the enriched functions of MCA while OTU143-*Fibrobacter* increased with acetate. These microbes either at genus or OTUs level had significant abundances in different regimes, however, they were not correlated with phonotypes very well. The reasons might be they were symbiotic with other microbiota. For example, *Treponema* does not utilize fiber, but it helps other bacteria to digest cellulosic materials (31).

The MCA-associated features, OTU87 (*Butyrivibrio hungatei*), OTU83 (*Prevotellamassilia timonensis*), OTU539 (*Abyssivirga alkaniphila*), and OUT93 (*Treponema pectinovorum*), correlated positively with NDF intake. Nevertheless, OTU539 associated with butyrate production and OTU93 related with propionate while OTU83 correlated with all 3 major VFA. The *Butyrivibrio hungatei* is the primary butyrate producers in the rumen and degraded effectively hemicellulose (32). *Prevotellamassilia timonensis* is a hemicellulose-degrading bacteria (33). *Abyssivirga alkaniphila* ferments saccharides, peptides and amino acids (34). *Fibrobacter* and *Treponema* synergetically break down the fiber components (35, 36). The function of OTU396 (*Pelobacter propionicus*) and OTU165 (*Marseilla massiliensis*) may be similar with OTU6. We observed their association with macronutrient and major VFA. Those MCA signatures cooperatively digest carbohydrate or protein and produce VFA. Increase of butyrate that is as an important regulator and stimulator of rumen development. Gorka et al. (37) revealed that the supplementation of alfalfa (NDF) could improve rumen development by increasing abundance of these synergistic bacteria. In addition, the OTU27 (*Prevotella falsenii*) might be nitrogen-specific bacteria since it correlated with CP intake and NH_3_-N very well though it was not classified as MCA signatures. A review reported some strains in *Prevotella* can degrade dietary proteins (31). Therefore, those signatures for regimes supplied with alfalfa contribute to both protein and carbohydrate utilization, and yield more nitrogen materials and VFA for host development. By contrast, the MRO signature microbiota cannot promote rumen functions at ruminant phase, however, they still may provide some baseline information of rumen in non-ruminant stage. Except *Sphingobacteriaceae*, OTU24 (*Cloacibacillus*) was another important specie for goats used only milk replacer as nutrient source. *Cloacibacillus* is a novel bacterium that degrades amino acids and produced VFAs (38). These MRO-associated signatures could be considered as passengers that contributed rumen development at specific time. Although little was known about their contribution of these bacteria, they were important for digestion of milk replacer and could be the primary strains impacted on late bacterial colonization.

The first limitation of this study is small sample size (6 per groups). However, it still showed good results between fluid diet and supplement of solid diet, providing some insights for future large scale studies. Second, the alfalfa in MCA groups were provide *ad libitum* resulting in less intake and less significant effects of adding alfalfa, but we did observed many bacteria related with fiber digestion due to significant fiber effects. Despite limitations, we confirm that signature microbiota for supplementary solid diet plays important roles in the promotion of rumen functions. Moreover, most of the MRO signatures function were not well described and are required for the identification of their functions by longitudinal measurements in further studies.

## Conclusions

Rumen fermentation and microbial composition were altered by supplementation of concentrate or concentrate+alfalfa, particularly the latte, in early life. The concentration of rumen VFA especially acetate, propionate and butyrate increased significantly when goats intake more nutrients from solid diet, and positively correlated with intake of crude protein, non-fiber carbohydrate and neutral detergent fiber. The membership and structure of rumen microbiota were altered. This study identified a set of signatures for supplementary solid diet regimes and validated their association with macronutrient intake and rumen fermentation. Also, it was the first time to use FishTaco for determination of link between those signatures’ abundances and rumen function shifts. Then, we performed network analysis to detect the interaction of signatures. By comprehensive integration, many members of bacteria having symbiotic relationship with signatures were classified. Therefore, for goat kids, extra nutrient from concentrate and/or forage manipulated core community structures by specific signature microbiota and their symbiotic partners, and then more volatile fatty acids were produced, and eventually rumen development and functions were promoted.

Our study answers several key questions in rumen microbiome affected by supplementary solid diet, and offers a foundation for studies aimed at improving ruminant health and production.

## Materials and Methods

### Goat kids, treatments and management

The experimental procedure was approved by the Chinese Academy of Agricultural Sciences Animal Ethics Committee, and humane animal care and handling procedures were followed throughout the experiment. This animal trial was conducted using Haimen goat kids at a commercial farm in the Jiangsu province, China.

A total of 72 Haimen goat kids (20 days old and average body weight 4.54± 0.51kg) were separated from their dams, and randomly allotted to three groups based on their following diets: milk replacer only (**MRO**), milk replacer + concentrate (**MRC**), milk replacer + concentrate + alfalfa pellets (**MCA**). Each group had six replicates and four kids per pen were as a replicate.

Goat kids remained with their mother and received breast milk from 0 to 20 days. During 20 to 60 days of age, they were separated with their dams and the above 3 kinds of diets were provided to corresponding groups. Other feeding management including vaccination, cleaning and disinfection of pens followed farm normal policy. All animals were fed with milk replacer from 20 to 60 days. Feeding amount of milk replacer were 2% body weight. Goat kids were fed 4 times a day (0600, 1200, 1800 and 2200) at 20-30 days, and thrice daily at 30-60 days (0600, 1200 and 1800). The milk replacer was dissolved with hot water cooled to 65-70 °C after boiling, and offered to goat kids when it was cooled to 40 ± 1 °C. The ratio of milk replacer to water was 1:6 (weight (g)/ volume (ml)). The milk replacer (China patent products ZL02128844.5) used in the experiment was provided by Beijing Precision Animal Nutrition Research Center. The concentrate with ingredients of corn, soybean etc. was purchased from Cargill Feed company, Nanjing. The alfalfa pellets purchased from Baofa Agriculture and Animal Husbandry Co. Ltd, Gansu, China had same diameter (4 mm) with concentrate diet. During the animal trial, all the goat kids had *ad libitum* access to water, the MRC and MCA kids were freely to access concentrate, and the MCA goats had extra free choice of alfalfa pellets. The nutritional levels of milk replacer, concentrate and alfalfa pellets are shown in Table S1.

### Sample collection and Chemical analysis

Daily feed intakes were recorded in animal trial. Feed samples were collected, dried in a forced-air oven at 65°C for 48 h and analyzed for crude protein (CP), non-fiber carbohydrate (NFC), and neutral detergent fiber (NDF) according to the Association of Official Analytical Chemists (39). Then, average daily intake of CP, NFC and NDF were calculated. Only data of table S1 (dietary composition) and table S2 (growth performance) were published in a Chinese journal paper (40), and other data, such as rumen fermentation parameters and microbiome analysis, were not published and used for the current draft.

Six goat kids (healthy and BW close to the average BW of the corresponding groups) from each group were selected and slaughtered for rumen samples collection. At 60 days of age, the goat kids were taken to an on-farm slaughterhouse, anesthetized using sodium pentobarbitone, and slaughtered by exsanguination from the jugular vein. Then, the rumen organs were taken out, and the ruminal content pH was measured immediately using pH electrode (PB-10; Sartorius, Goettingen, Germany). Around 10 ml rumen content were sampled from the whole mixed rumen digesta and stored at −80°C for sequencing. The rumen fluid around 10 ml filtered through four layers of gauze was placed in a 15 ml centrifuge tube immediately frozen at −20°C for measurement of rumen fermentation. Determination of rumen fluid NH_3_-N concentration by phenol-sodium hypochlorite colorimetric method after the liquid was thawed at 4°C. The microbial proteins were analyzed according to the method described by (41). The volatile fatty acids (VFA) in rumen fluid were quantified by gas chromatography (42) using methyl valerate as internal standard in an Agilent 6890 series GC equipped with a capillary column (HP-FFAP19095F-123, 30 m, 0.53 mm diameter and 1 mm thickness).

### DNA extraction and 16S rRNA gene sequencing

Rumen fluid samples were thawed on ice and microbial DNA was extracted using a commercial DNA Kit (Omega Bio-tek, Norcross, GA, U.S.) according to manufacturer’s instructions. Total DNA quality analysis using a Thermo NanoDrop 2000 UV spectrophotometer and 1% agarose gel electrophoresis. The V3-V4 region of the bacteria 16S ribosomal RNA genes were amplified by PCR (95 °C for 3 min, followed by 30 cycles at 98°C for 20 s, 58 °C for 15s, and 72 °C for 20 s and a final extension at 72 °C for 5 min) using indexes and adaptors-linked universal primers (431 F:ACTCCTACGGGRSGCAGCAG; 806R: GGACTACVVGGGTATCTAATC). PCR reactions were performed in 30 μL mixture containing 15 μL of 2 × KAPA Library Amplification Ready Mix, 1 μL of each primer (10 μM), 50ng of template DNA and ddH_2_O. All PCR products were normalized and quantified by a Qubit 2.0 Fluorometer (Thermo Fisher Scientific, Waltham, US). Amplicon libraries were mixed using all qualified products and sequenced with Illumina HiSeq PE250 plateform at Realbio Technology Genomics Institute (Shanghai, China).

### Sequencing Data Processing

Raw sequences were filtered through a quality control pipeline using the Quantitative Insight into Microbial Ecology (QIIME) tool kit (43). The chimeras and singletons were detected and removed by Usearch software, and the high quality sequences were clustered into operational taxonomic units (OTUs) at the 97% similarity level. Samples were normalized to 24136 sequencing reads. The representative sequence was classified based on the Ribosomal Database Project (RDP) database (44) at the default confidence threshold of 0.8, trained on the SILVA reference database (release 111) (45). The alpha diversities (Shannon Index and Observed species), and beta diversity (Unweighted and Weighted Unifrac distance) were calculated. The ANalysis Of SIMilarity (ANOSIM) test was used to examine the statistical significance of differences in beta diversity. The datasets in the current study are available in the NCBI BioProject database with the BioProject ID PRJNA544381 (https://www.ncbi.nlm.nih.gov/sra/PRJNA544381).

### Data Analysis

Rumen fermentation parameters were shown using bar charts made in R (v3.6.0) by ‘ggplot2’ package. The Anova test was used for significance calculation after detection of homogeneity of variance. After the global test was significant, a post-hoc analysis (Tukey’s HSD test) was performed to determine which group of the independent variable differ from each other group.

Alpha diversity of the rumen microbial data among three treatments was tested using Kruskal–Wallis test and a post-hoc Dunn Kruskal-Wallis multiple comparison with Bonferroni adjustment to evaluate differences between two groups, and boxplots were made in R (‘ggpubr’ packages). Beta diversity was visualized with PCoA plot through R.

RandomForest classification model was performed to identify the top microbiome signatures to differentiate 3 supplementary feeding regimes. R package ‘AUCRF’ (v.1.1) was used to process RandomForest model and select optimal variables based on the area-under-the receiver operator characteristic curve (AUC) of the RandomForest method (AUCRF) (46). The relative abundances of all the microbiota were included for predictors selection. The ‘ntree’ parameters was set at 10,000 in the model. For calculation the probability of each selected variable, a 10-fold cross validation analysis and 20 times repetitions of cross validation were performed. The model accuracy, including AUC, sensitivity and specificity of variables, was calculated using the ‘pROC’ package (v.1.13). Thus, variables importance plot was generated based on the importance scores (Mean Decrease in Accuracy, MDA) of optimal features and their boxplots of selected features were drawn in R.

RandomForest regression model was used to select the rumen microbiota that were important for average daily intake of major nutrients (i.e., CP, NDF and NFC) and rumen fermentation parameters. The model was run in R software using ‘RandomForest’ package (v 4.6-14). The percent variance explained was reported for the estimation of accuracy of regression model. The top 50 selected features were then analyzed Pearson correlation with those macro indicators respectively.

Predictive function analysis was performed using the PICRUSt algorithm based on the Kyoto Encyclopedia of Genes and Genomes (KEGG) classification using the closed-reference OTUs (47). The Functional Shifts’ Taxonomic Contributors (FishTaco) software was used to find the rumen bacteria driving the functional shifts between supplementary regimes in this study. A taxonomic abundance at OTUs’ level and functional abundance profile at levels 3 from the PICRUSt analysis were used. In pairwise comparisons, we labeled MRO groups as control and MRC or MCA as case, and tested MRC as control vs MCA as case. Each functional shift was grouped into case-associated with driving case-enrichment or attenuating case-enrichment, and control-associated driving case-enrichment or attenuating case-enrichment. The output results visualization was performed in FishTacoPlot package in R (Version 3.6.0).

Network analysis was performed by calculating all possible Pearson rank correlation coefficients (ρ) between microbial pairs. To minimize the occurrence of spurious associations, we considered a valid co-occurrence between two different taxa if a correlation co-efficiency over 0.6 or less than 0.6 and statistically significant. The network was demonstrated by using the ‘igraph’ package in R with edges connecting nodes (bacterial taxa). The subnetworks in regimes were produced based on the betweenness cluster calculated by the Girvan-Newman algorithm (48).

## Acknowledgements

This study was funded by grants from National Key R&D Program Projects(2018YFD0501902), National Natural Science Foundation of China (31872385) and National Technical System Construction of Mutton Sheep Industry(CARS-39).

## Supplemental material

Supplementary information is available at the ISME journal’s website.

